# Forward genetic screens identify mechanisms of resistance to small molecule lactate dehydrogenase inhibitors

**DOI:** 10.1101/2023.09.30.560315

**Authors:** Anderson R Frank, Florentina Vandiver, David G McFadden

**Affiliations:** Department of Internal Medicine, Division of Endocrinology, University of Texas Southwestern Medical Center, Dallas, TX 75390, USA; Department of Biochemistry, University of Texas Southwestern Medical Center, Dallas, TX 75390, USA; Harold C. Simmons Comprehensive Cancer Center, University of Texas Southwestern Medical Center, Dallas, TX 75390, USA; Program in Molecular Medicine, University of Texas Southwestern Medical Center, Dallas, TX 75390, USA

## Abstract

Altered metabolism is a hallmark of cancer; however, it has been difficult to specifically target metabolism in cancer for therapeutic benefit. Cancers with genetically defined defects in metabolic enzymes constitute a subset of cancers where targeting metabolism is potentially accessible. Hürthle cell carcinoma of the thyroid (HTC) tumors frequently harbor deleterious mitochondrial DNA (mtDNA) mutations in subunits of complex I of the mitochondrial electron transport chain (ETC). Previous work has shown that HTC models with deleterious mtDNA mutations exhibit mitochondrial ETC defects that expose lactate dehydrogenase (LDH) as a therapeutic vulnerability. Here, we performed forward genetic screens to identify mechanisms of resistance to small molecule LDH inhibitors. We identified two distinct mechanisms of resistance: upregulation of an LDH isoform and a compound-specific resistance mutation. Using these tools, we demonstrate that the anti-cancer activity of LDH inhibitors in cell line and xenograft models of complex I-mutant HTC is through on-target LDH inhibition.

## INTRODUCTION

Metabolism supports cell homeostasis, growth, and division by extracting energy from nutrients and generating important biochemical intermediates and signaling molecules (1). Early work by Warburg suggested that cancer cells have alterations in their metabolism that enable oncogenesis, and an ever-growing body of evidence has reinforced that altered metabolism supports cancer cell growth (2–5). The early success of anti-metabolite chemotherapies demonstrated that metabolism is a therapeutically tractable cellular process in cancer; however, it could be argued that these therapies do not specifically target altered metabolism (6, 7). More recently, the discovery that gain-of-function mutations in the tricarboxylic acid (TCA) cycle enzymes, isocitrate dehydrogenase 1 and 2 (IDH1 and IDH2), promote oncogenesis through the production of (R)-2-hydroxyglutarate has enabled the development of clinically approved small molecule drugs that specifically inhibit mutant IDH (8–14). Despite these successes, the ability to target altered metabolism in cancer remains limited, likely due to the inherently adaptive nature of cellular metabolism.

A central component of cellular metabolism is the mitochondrial electron transport chain (ETC) which, among other activities, generates adenosine triphosphate (ATP) from reduced electron carriers generated in the TCA cycle (15, 16). Under conditions where mitochondrial ETC activity is impaired, cells rely on the combined actions of glycolysis and fermentation to generate the ATP required for cell survival. The mitochondrial ETC is unique in that the protein subunits that form ETC complexes are encoded by both the nuclear and mitochondrial genomes. Sequencing studies have found that while somatic mitochondrial DNA (mtDNA) mutations are common in cancer, these mtDNA mutations are typically not loss-of-function, consistent with the need for a functional mitochondrial ETC to support tumor growth (17–20). While most tumors do not enrich for loss-of-function mtDNA mutations, approximately 10% of kidney, colon, and thyroid tumors harbor high allelic fraction, truncating mtDNA mutations (21, 22). Additionally, oncocytic tumors (oncocytomas) – a class of tumors characterized by significant mitochondrial accumulation – frequently harbor deleterious high allelic fraction mtDNA mutations (23–26).

We and others have found that approximately 60% of Hürthle cell carcinoma of the thyroid (HTC, also known as oncocytic thyroid carcinoma) tumors harbor near-homoplasmic missense and loss-of-function mtDNA mutations affecting complex I of the mitochondrial ETC (23, 25–28). Recent work has demonstrated that HTC models harboring these high allelic fraction mtDNA mutations exhibit severe mitochondrial ETC defects that impair central carbon metabolism (28, 29). These genetically encoded respiratory defects expose diverse metabolic processes, including glycolysis and fermentation, that have the potential to be targeted therapeutically (28, 29). Whether these findings are applicable in other colon, kidney, or thyroid tumors with disruptive mtDNA mutations remains to be seen; however, our work raises the possibility that altered metabolism could be targeted in these cancers with small molecule lactate dehydrogenase (LDH) inhibitors (21, 22, 29).

While targeted therapies have provided significant advances in the treatment of cancer, resistance to targeted therapies is a common occurrence (30–33). Diverse mechanisms for resistance have been identified including upregulation of the cellular target (33, 34), mutations in drug-binding sites (33–36), and even transition to a different cell state (32, 36). Understanding these resistance mechanisms can inform treatment regimens and lead to the development of next-generation therapeutics (30, 33, 37). Attempts to identify resistance mechanisms in cultured cancer cells have previously utilized long-term culture with sub-lethal concentrations of a compound of interest; however, these approaches might bias towards non-genetic mechanisms of resistance (i.e., upregulation of drug efflux pumps). Several groups have utilized the DNA mismatch repair (MMR)-deficient colorectal cancer cell line, HCT116, to identify mutation-based resistance mechanisms (38–41). The ability to identify these mechanisms in HCT116 is afforded by the increased mutagenic rate found in these cells due to MMR defects. Recently, we and others have demonstrated that this approach can be successfully adopted in MMR-proficient cell lines through CRISPR-Cas9-mediated silencing of MMR genes or inducible depletion of MMR proteins (42, 43).

In this study, we used CRISPR-Cas9 to generate an MMR-deficient HTC cell line and performed forward genetic screens to identify mechanisms of resistance to LDH inhibitors (LDHi). Through these screens we identified two LDHi resistance mechanisms: i) upregulation of *LDHB*, which induces pan-LDHi resistance, and ii) a mutation in *LDHA* that confers compound-specific resistance to the LDH inhibitor, NCGC00420737.

## RESULTS

### MSH2 loss enables forward genetic screening in HTC cells

To facilitate forward genetic screening for LDH inhibitor resistance mechanisms in HTC, we utilized an HTC cell line model, NCI-237^UTSW^, that we previously found to be sensitive to LDH inhibition (29). We used CRISPR-Cas9 gene editing to generate isogenic MSH2-wild-type or MSH2-null populations of NCI-237^UTSW^ cells, a strategy that we have employed to enable forward genetic screening in otherwise MMR-proficient cell lines (Figures 1A and S1A) (42, 43). As a proof-of-concept, we tested whether MSH2 loss would enable the identification of cells with specific resistance to the NEDD8 activating enzyme (NAE) inhibitor, MLN4924 (44). Prior studies have identified a series of mutations that confer specific resistance to MLN4924, thus, we considered this a suitable test compound (40–43, 45).

**Figure 1:**
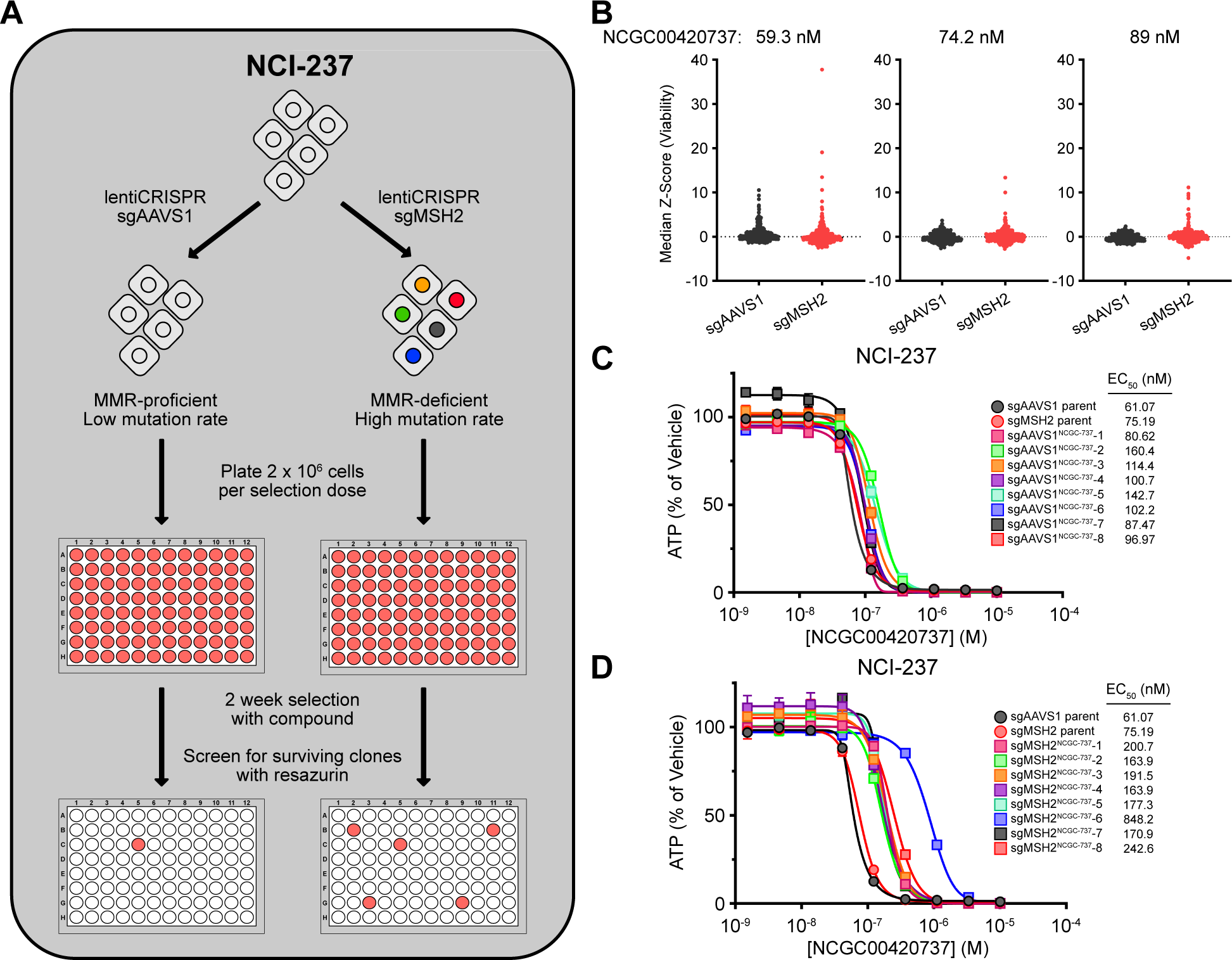
MSH2 loss enables forward genetic screens for compound resistance in NCI-237^UTSW^. A) Schematic for forward genetic screening in NCI-237^UTSW^. B) Resazurin viability assay for NCI-237^UTSW^ cells treated with indicated concentrations of NCGC00420737 for 14 days; *n* = 384 wells/cell line/condition. Outliers (positive Z-score) indicate viable clones. C) Viability assay for NCI-237^UTSW^ sgAAVS1 clones treated with NCGC00420737 for 3 days; *n* = 2 replicates. D) Viability assay for NCI-237^UTSW^ sgMSH2 clones treated with NCGC00420737 for 3 days; *n* = 2 replicates. Data are plotted as mean ± SEM of the indicated number of replicates. The data displayed in 1C and 1D were generated during the same experiment and the data for sgAAVS1 and sgMSH2 parent in 1C and 1D are the same data.

We subjected MSH2-null and MSH2-wild-type NCI-237^UTSW^ cells to selection with a lethal dose of MLN4924 and observed significantly more viable cells following treatment with 1.32 μM MLN4924 in the MSH2-null population (Figures S1B and S1C). Surviving clones from both the MSH2-null and MSH2-wild-type populations were expanded and sensitivity to MLN4924 was assessed by compound dose-response assay. All MSH2-null clones tested (6/6) were resistant to MLN4924, with an average EC_50_ increase of 5.53-fold (range 3.88- to 7.58-fold) relative to the MSH2-wild-type and MSH2-null parental populations (Figure S1D). Additionally, these clones were not cross-resistant to the LDHA/LDHB inhibitor, NCGC00420737, suggesting the resistance mechanism was compound-specific (Figure S1E). Clones recovered from the MSH2-wild-type selection exhibited modest resistance as evidenced by a 1.75-fold EC_50_ increase (range 0.66- to 2.50-fold) (Figures S1F and S1G). Targeted sequencing of *UBA3* in the MSH2-null resistant clones identified a recurrent heterozygous missense mutation (Chr3:69056662T>C; p.Tyr352His) causing a Tyr-to-His substitution at amino acid position 352 (Figures S1H and S1I). This mutation was previously reported to cause resistance to MLN4924 treatment (43, 45). These data suggested that MSH2 loss in NCI-237^UTSW^ cells induced a mutagenic state amenable to forward genetic screens for compound resistance mechanisms.

### MMR deficiency supports the identification of LDHi-resistant cells

We next asked whether resistance mechanisms to the pyrazole-based LDHA/LDHB inhibitor, NCGC00420737 (46, 47). could be identified through forward genetic screens. MSH2-null and MSH2-wild-type NCI-237^UTSW^ were exposed to 59.3, 74.2, or 89 nM NCGC00420737 for two weeks before screening for surviving clones (Figures 1B and S2A). MSH2-null cells gave rise to more resistant clones than MSH2-wild-type cells, suggesting that survival following NCGC00420737 treatment was mutation-driven (Figure 1B). We expanded and assayed sensitivity to NCGC00420737 from MSH2-wild-type and MSH2-null clones raised against 59.3 nM NCGC00420737, as well as MSH2-null clones raised against 74.2 and 89 nM NCGC00420737 (no MSH2-wild-type clones were recovered from these selections). MSH2-null, but not MSH2-wild-type clones from the 59.3 nM NCGC00420737 selection displayed resistance to NCGC00420737, with an average EC_50_ increase of 3.96-fold (range 2.41- to 12.45-fold) (Figures 1C and 1D). Clones recovered from the 74.2 nM and 89 nM NCGC00420737 selections displayed EC_50_ increases of 9.53- and 15.44-fold, respectively (Figure S2D). NCGC00420737-resistant clones were not cross-resistant to MLN4924 or the proteasome inhibitor bortezomib, suggesting compound-specific resistance (Figures S2B, S2C, and S2E).

### LDHB upregulation is a pan-LDHi resistance mechanism

In addition to NCGC00420737, we previously found that NCI-237^UTSW^ cells were sensitive to a chemically distinct LDHA/LDHB inhibitor, (R)-GNE-140 (29). As NCGC00420737 and (R)-GNE-140 are both competitive LDHA/LDHB inhibitors (46–48), we tested whether NCGC00420737-resistant clones display cross-resistance to (R)-GNE-140. We performed (R)-GNE-140 compound dose-response assays with NCGC00420737-resistant clones and found that 7/10 clones displayed cross-resistance to (R)-GNE-140 with an average 4.42-fold increase in (R)-GNE-140 EC_50_ (range 1.45- to 7.92-fold) (Figures 2A, 2B, and S3A). These results highlighted that there were two distinct resistance profiles among our isolated clones: i) clones that displayed greater resistance to NCGC00420737 (approximately 10-fold) with minor resistance to (R)-GNE-140 (less than 2-fold), and ii) clones with pan-LDHi resistance, displaying moderate resistance to both NCGC00420737 (approximately 3-fold) and (R)-GNE-140 (approximately 5.5-fold).

**Figure 2:**
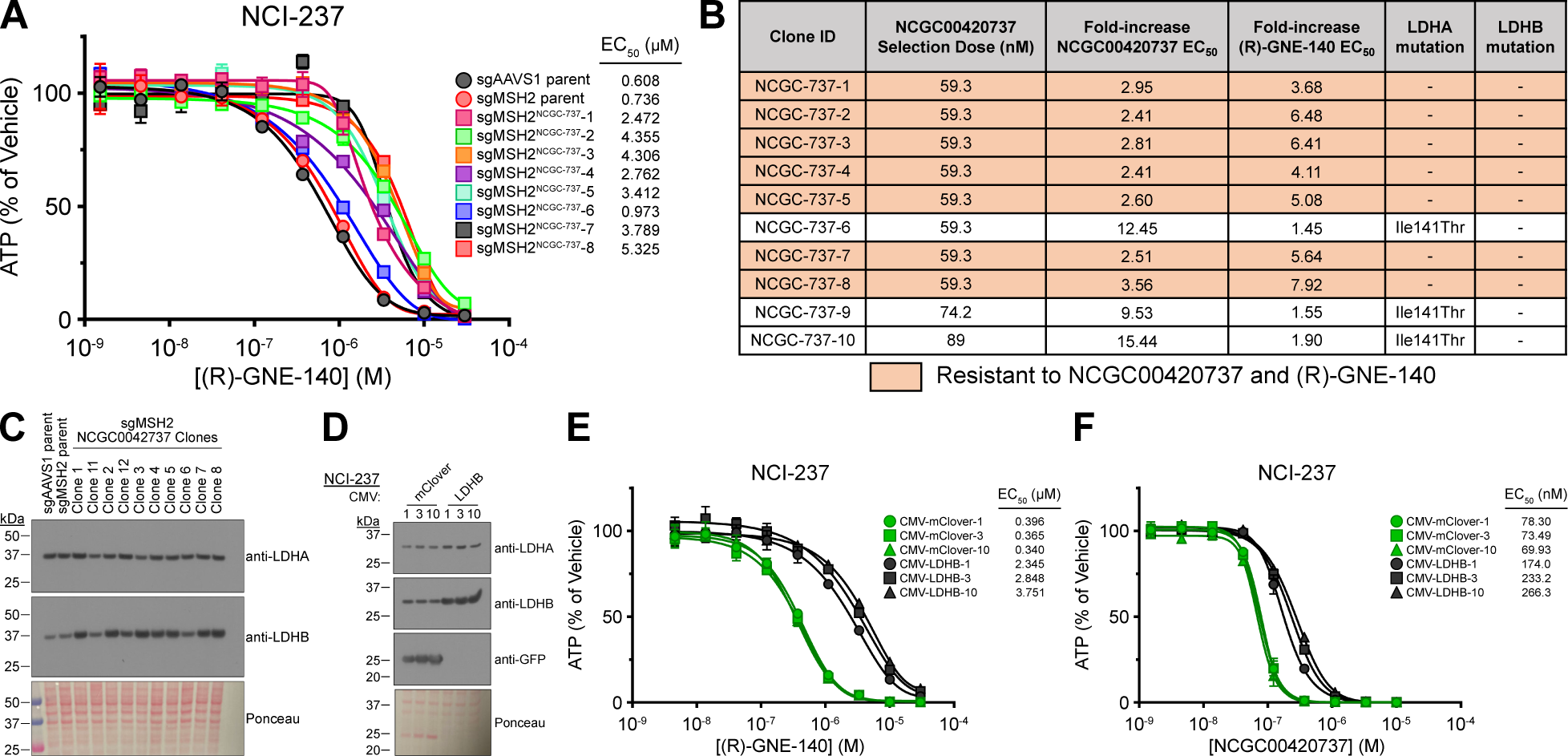
LDHB overexpression drives resistance to multiple LDH inhibitors. A) Viability assay for NCI-237^UTSW^ sgMSH2 clones treated with (R)-GNE-140 for 3 days; *n* = 2 replicates. B) Table of fold-change in EC_50_ values and *LDHA* or *LDHB* mutation status for NCI-237^UTSW^ sgMSH2 NCGC00420737-resistant clones. C) Immunoblot for LDHA and LDHB in NCI-237^UTSW^ sgMSH2 NCGC00420737-resistant clones. D) Immunoblot for LDHA, LDHB, and GFP in NCI-237^UTSW^ cells expressing mClover or LDHB. E) Viability assay for NCI-237^UTSW^ cells expressing indicated constructs treated with (R)-GNE-140 for 3 days; *n* = 2 replicates. F) Viability assay for NCI-237^UTSW^ cells expressing indicated constructs treated with NCGC00420737 for 3 days; *n* = 2 replicates. Data are plotted as mean ± SEM of the indicated number of replicates.

Given that our resistant clones were isolated in a mutation-dependent manner, we performed targeted sequencing of *LDHA* and *LDHB* to identify potential resistance mutations. We identified a single heterozygous missense mutation in *LDHA* in clones that displayed resistance to NCGC00420737, but not (R)-GNE-140 (NCI-237^UTSW^ sgMSH2^NCGC-737^ Clones 6, 9, and 10). This mutation (Chr11:18402846T>C; p.Ile141Thr) causes an Ile-to-Thr substitution at amino acid position 141 in LDHA (Figures 2B and S3B). We did not, however, identify any clonal, nonsynonymous mutations in the *LDHA* or *LDHB* coding sequence of pan-LDHi resistant clones.

In the absence of a protein-altering mutation, what mechanisms could drive resistance to multiple inhibitors of the same target? Previous studies have demonstrated that upregulation of a compound’s cellular target can drive resistance (32, 34), therefore, we asked whether upregulation of *LDHA* or *LDHB* coincided with pan-LDHi resistance. We found that pan-LDHi resistant clones displayed higher levels of LDHB, but not LDHA, and that this was driven by upregulation of *LDHB* transcripts (Figures 2C and S3C). These data suggested that overexpression of LDHB would cause resistance to both NCGC00420737 and (R)-GNE-140. To test whether LDHB upregulation was sufficient to confer pan-LDHi resistance, we generated NCI-237^UTSW^ and TPC-1 cells with lentiviral expression of mClover or LDHB (Figures 2D and S3D). Consistent with our hypothesis, cells overexpressing LDHB were resistant to (R)-GNE-140 and NCGC00420737 (Figures 2E, 2F, S3E, and S3F). Combined, the above data demonstrate that LDHB upregulation is sufficient to confer resistance to multiple LDH inhibitors.

### LDHA^I141T^ encodes resistance to NCGC00420737

Through sequencing of *LDHA* and *LDHB* in NCGC00420737-resistant clones, we identified a recurrent missense mutation in LDHA – LDHA^I141T^ (Figures 2B and S3B). Mapping the location of Ile141 in a previously determined co-crystal structure of LDHA and NCGC00420737 revealed that this amino acid lies within a binding pocket that is contacted by the sulfonamide group of NCGC00420737 (Figure 3A) (47). This structural data suggested that mutation of Ile141 might disrupt the interaction between LDHA and NCGC00420737, thus providing a potential explanation for why this mutation would cause resistance to NCG00420737.

**Figure 3:**
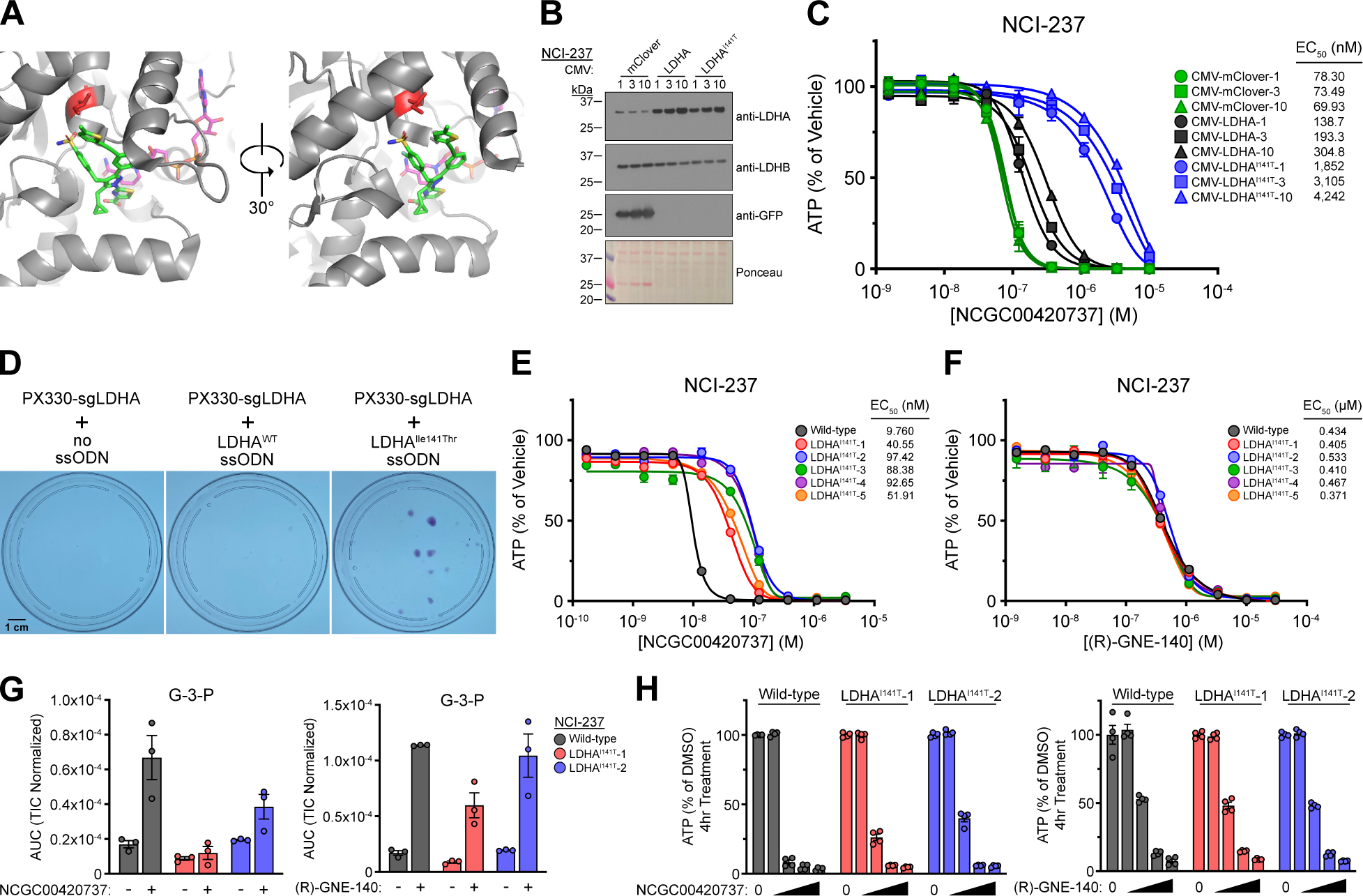
LDHA^I141T^ encodes resistance to NCGC00420737. A) Co-crystal structure of LDHA and NCGC00420737. LDHA is colored in gray; Ile141 is colored in red; NAD^+^ is colored in magenta; NCGC00420737 is colored in green. PDB: 6Q13. B) Immunoblot for LDHA, LDHB, and GFP in NCI-237^UTSW^ cells expressing mClover, LDHA, or LDHA^I141T^. C) Viability assay for NCI-237^UTSW^ cells expressing indicated constructs treated with NCGC00420737 for 3 days; *n* = 2 replicates. D) Crystal violet staining of NCI-237^UTSW^ cells transfected with indicated plasmids and treated with 333 nM NCGC00420737 for 2 weeks. E) Viability assay for NCI-237^UTSW^ cells with endogenous LDHA^I141T^ treated with NCGC00420737 for 3 days; *n* = 2 replicates. F) Viability assay for NCI-237^UTSW^ cells with endogenous LDHA^I141T^ treated with (R)-GNE-140 for 3 days; *n* = 2 replicates. G) G-3-P levels in indicated NCI-237^UTSW^ cells treated with 40 nM NCGC00420737 (left) or 1 µM (R)-GNE-140 (right) for 4 hours; *n* = 3 replicates. H) ATP levels in indicated indicated NCI-237^UTSW^ cells treated with 0, 0.014, 0.123, 1.11, or 10 µM NCGC00420737 (left) or 0, 0.041, 0.370, 3.33, or 30 µM (R)-GNE-140 (right) for 4 hours; *n* = 3 replicates. Data are plotted as mean ± SEM of the indicated number of replicates. G-3-P = glyceraldehyde-3-phosphate.

We tested whether LDHA^I141T^ encodes resistance to NCGC00420737 by generating NCI-237^UTSW^ and TPC-1 cells with ectopic expression of mClover, LDHA, or LDHA^I141T^ and subjecting these cells to LDHi dose-response assays (Figures 3B, 3C, S4A, and S4B). Cells expressing LDHA displayed a modest increase in NCGC00420737 EC_50_ (NCI-237^UTSW^: 1.9- to 4.1-fold increase; TPC-1: 1.7- to 2.2-fold increase) while cells expressing LDHA^I141T^ were significantly more resistant to NCGC00420737 (NCI-237^UTSW^: 25.1- to 57.4-fold EC_50_ increase; TPC-1: 9.9- to 14.2-fold EC_50_ increase) (Figures 3C and S4B). Consistent with the results we observed for clones isolated from our screen that harbored LDHA^I141T^ mutations, overexpression of LDHA or LDHA^I141T^ did not confer resistance to (R)-GNE-140 (Figures 2A, S3A, S4C, and S4D). Why does LDHA^I141T^ encode resistance to NCGC00420737 but not (R)-GNE-140? Analysis of a previously determined co-crystal structure of LDHA and (R)-GNE-140 revealed that (R)-GNE-140 does not bind LDHA in the vicinity of Ile141, providing an explanation for the compound-specific effects we observed for the LDHA^I141T^ mutation (Figure S4E) (48).

We next used CRISPR-Cas9 genome editing to assess whether installing the LDHA^I141T^ mutation at the endogenous *LDHA* locus would confer resistance to NCGC00420737. Introduction of Cas9, an sgRNA targeting *LDHA*, and a single-stranded oligodeoxynucleotide repair template encoding LDHA^I141T^ promoted NCI-237^UTSW^ cell survival following NCGC00420737 treatment (Figure 3D). Multiple subclones recovered from the LDHA^I141T^ knock-in condition harbored the LDHA^I141T^ mutation and displayed resistance to NCGC00420737, but not (R)-GNE-140 or toxins with distinct mechanisms-of-action (Figures 3E, 3F, S4F, and S4G). Interestingly, the fold-increase in EC_50_ (approximately 5- to 10-fold) observed in cells with LDHA^I141T^ introduced at the endogenous locus was more similar to clones isolated from our screen with heterozygous LDHA^I141T^ mutations than cells with overexpression of LDHA^I141T^ (Figures 1D, 2B, 3C, and 3E).

To determine whether the rescue effect of LDHA^I141T^ was through impaired NCGC00420737 action in cells, we analyzed the metabolic response to LDHi treatment in wild-type NCI-237^UTSW^ cells and two independent LDHA^I141T^ clones. Cells were treated with NCGC00420737 or (R)-GNE-140 for four hours and intracellular metabolite levels were measured using gas chromatography-mass spectrometry (GC-MS). Wild-type NCI-237^UTSW^ cells treated with NCGC00420737 exhibited an accumulation of the upper glycolytic intermediate glyceraldehyde-3-phosphate (G-3-P) and its isomer, dihydroxyacetone phosphate (DHAP), and this effect was blunted in cells harboring the LDHA^I141T^ mutation (Figure 3E and S4H). Consistent with specific rescue of NCGC00420737, both wild-type and LDHA^I141T^-mutant cells exhibited an increase in G-3-P and DHAP levels following (R)-GNE-140 treatment (Figure 3G and S4H). Additionally, we measured ATP levels in NCI-237^UTSW^ wild-type and LDHA^I141T^ cells following short-term treatment with NCGC00420737 and observed that ATP levels were buffered in LDHA-mutant cells; however, (R)-GNE-140 treatment caused a similar decrease in ATP levels in both wild-type and LDHA^I141T^ cells (Figure 3G). These results demonstrate that the *in vitro* toxicity of NCGC00420737 is through on-target LDH inhibition.

### Anti-tumor activity of NCGC00420737 in an HTC xenograft model is due to on-target LDH inhibition

Recent work has highlighted how small molecules identified through *in vitro* biochemical screens can exert their anti-cancer activities through a different mechanism than what was originally ascribed to the molecule (i.e., the cellular target responsible for cell death is different than the original screen target) (49–52). Additionally, inhibitors of metabolic processes can have impaired efficacy *in vivo* compared to *in vitro* due to differences in the extracellular milieu and the resulting alterations in cellular metabolism (53, 54). Thus, we utilized the LDHA^I141T^ mutation as a genetic tool to assess whether NCGC00420737 efficacy against complex I-mutant HTC *in vivo* was due to on-target LDH inhibition.

Wild-type and LDHA^I141T^ mutant NCI-237^UTSW^ subcutaneous xenografts were formed in immunodeficient mice and tumor-bearing mice received 60 mg/kg NCGC00420737 intravenously on a ‘5 days on, 2 days off’ schedule. We observed that 60 mg/kg NCGC00420737 caused a significant anti-tumor response in wild-type NCI-237^UTSW^ xenografts; however, NCI-237^LDHA-I141T^ xenografts grew at approximately the same rate in vehicle and treatment conditions (Figure 4A, 4B, and S5A). Combined, the above data demonstrate that the *in vivo* efficacy of NCGC00402737 in a model of complex I-mutant HTC is through on-target LDH inhibition and that LDHA^I141T^ provides a genetic tool for assessing on-target NCGC00420737 activity in cultured cells and animals.

**Figure 4:**
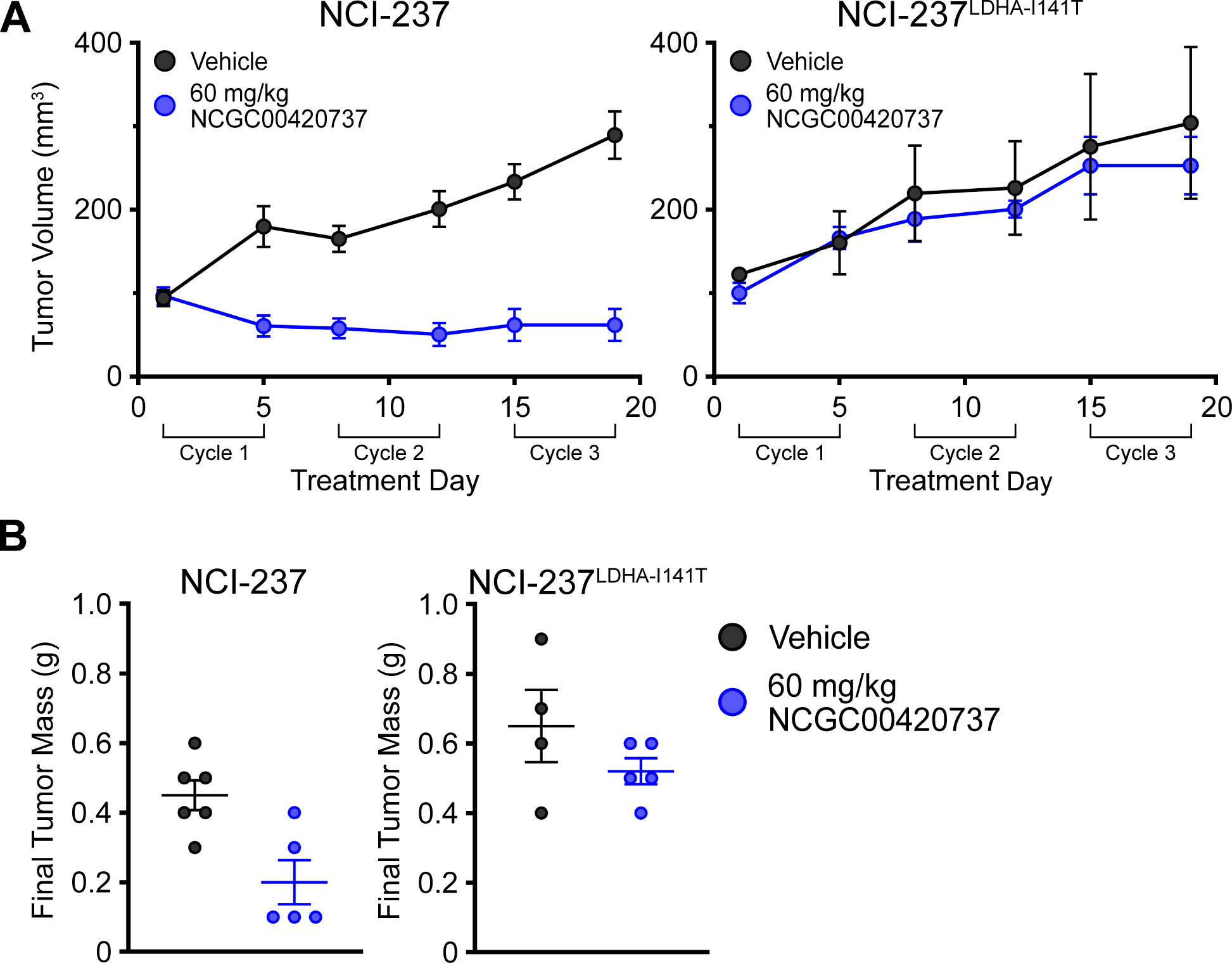
Anti-tumor activity of NCGC00420737 in a xenograft model of HTC is due to on-target LDH inhibition. A) Tumor volumes for NCI-237^UTSW^ xenografts in animals receiving vehicle (*n* = 6) or 60 mg/kg NCGC00420737 (*n* = 5) (Left). Tumor volumes for NCI-237^UTSW^ ^LDHA-I141T^ xenografts in animals receiving vehicle (*n* = 4) or 60 mg/kg NCGC00420737 (*n* = 5) (Left). Vehicle or compound was administered once daily via jugular vein catheter. A final compound administration was performed, and animals were sacrificed 1 hour after receiving compound. B) Final tumor mass for NCI-237^UTSW^ (left) and NCI-237^UTSW^ ^LDHA-I141T^ (right) xenografts from A). Data are plotted as mean ± SEM of the indicated number of replicates.

## DISCUSSION

Acquired resistance to targeted anti-cancer therapies can provide clues about a drug’s mechanism-of-action and inform the development of next-generation therapeutics. Here, we used forward genetic screening to identify two mechanisms of resistance to LDH inhibition: i) a compound-resistant allele specific for NCGC00420737, and ii) upregulation of *LDHB*.

Using LDHA^I141T^ as a compound-specific genetic tool, we confirmed that the cytotoxic and anti-tumor effects of NCGC00420737 treatment in a model of complex I-mutant HTC are due to on-target LDH inhibition. While NCI-237^UTSW^ cells and tumors harboring the LDHA^I141T^ mutation displayed resistance to NCGC00420737 in growth assays and animal dosing studies, the short-term metabolic rescue afforded by LDHA^I141T^ was less pronounced. This suggests that even relatively minor interventions that raise the levels of metabolites such as ATP could rescue viability in complex I-mutant HTC cells subjected to LDH inhibition. While NCGC00420737 and other LDH inhibitors have not been approved for clinical use, the LDHA^I141T^ mutation could be a useful genetic tool in the clinical development of NCGC00420737. Previous work has highlighted that hemolysis is a potential dose-limiting toxicity of NCGC00420737 (29, 55) and the generation of a genetically-engineered mouse model with LDHA^I141T^ (assuming this mutation is tolerated at the organismal level) could aid in assessing the on-target toxicity of NCGC00420737. Additionally, LDHA^I141T^ could be utilized as a primary screening target, or a counter-screening tool, in high-throughput chemical screens to identify next-generation LDH inhibitors.

Regarding the mutational space that confers resistance to NCGC00420737 and other competitive LDH inhibitors, we only recovered a single missense mutation, and this mutation was rare (approximately 1 in 2,000,000 cells harbored LDHA^I141T^ based on the number of recovered clones with this mutation). These results suggest that Ile141 is one of the few amino acid residues in LDHA where a mutation that both preserves enzyme activity while blocking compound activity can be achieved. Given that NCGC00420737 is a competitive inhibitor, this is not surprising as active site residues that are required for enzyme function also coordinate compound binding. By contrast, greater mutational diversity can be seen with allosteric inhibitors or other compounds that do not bind enzyme active sites (38, 56). Another possibility is that the mutational spectrum generated by MSH2 loss and MMR deficiency, which favors C>T and T>C substitutions (https://cancer.sanger.ac.uk/signatures/sbs/) (57, 58), does not fully capture the mutation space that could drive resistance. Targeted mutagenesis methods such as CRISPR-Cas9 base editing screens or saturation mutagenesis screens could expand the number of known LDHi compound-resistant alleles (56, 59, 60).

Additionally, our results suggest that *LDHB* upregulation is likely to be a more common resistance mechanism than compound-resistant mutations. Beyond conferring resistance to diverse LDH inhibitors, upregulation of *LDHB* could increase cytosolic NAD^+^ levels and subsequently increase the glycolytic capacity of cells by providing substrate for glyceraldehyde-3-phosphate dehydrogenase (GAPDH). Thus, *LDHB* overexpressing clones are likely more frequent within the population as *LDHB* upregulation represents a resistance mechanism where there is minimal trade-off or disadvantage to the genetic events that promote resistance. While the underlying cause of *LDHB* upregulation remains unknown, *LDHB* upregulation appears to be a mutation-driven process, as we only isolated LDHi-resistant clones from selections in MSH2-null NCI-237^UTSW^ cells (Figures 1C and 1D). Potential explanations include mutations in the 5’ or 3’ untranslated regions of the *LDHB* transcript that contribute to increased mRNA stability, or amplification of chromosome 12, which harbors the *LDHB* locus. Mutations in gene regulatory regions or protein-altering mutations in transcriptional regulators could also contribute to *LDHB* upregulation. These possibilities warrant additional studies to uncover the genetic underpinnings of *LDHB* transcriptional induction in LDHi-resistant clones with the possibility to expand upon known mechanisms that drive drug resistance.

## MATERIALS AND METHODS

### Human cell lines

The generation of NCI-237^UTSW^ (male) was previously described (29). TPC-1 (female) was obtained from Dr. Sareh Parangi (Massachusetts General Hospital, Boston, MA, USA). HEK 293T/17 (female) were purchased from ATCC (CRL-11268). All cell lines were subjected to short-tandem repeat (STR) profiling performed by the Eugene McDermott Center for Human Growth and Development Sequencing Core. Cell lines were periodically tested for mycoplasma contamination using a PCR-based assay.

### Cell culture

Cell lines were maintained at 37°C with 5% CO_2_. NCI-237^UTSW^ were cultured in either DMEM (Sigma D6429) supplemented with 10% FBS, 2 mM L-glutamine, 1% penicillin/streptomycin, and 50 µg/mL uridine or human plasma-like media (HPLM) (54) supplemented with 2% FBS, 1% penicillin/streptomycin, and 1X insulin-transferrin-selenium. HPLM pool stocks were prepared and stored as previously described (54) and HPLM was assembled by combining pool stock solutions in deionized water, adjusting pH to ∼7.2 with hydrochloric acid, and filtering through a 0.22 µm PES filter (Corning 431097). Prepared HPLM was used within 3-4 days and supplements were added daily before use. TPC-1 and HEK 293T/17 were cultured in DMEM (Sigma D6429) supplemented with 10% FBS, 2 mM L-glutamine, 1% penicillin/streptomycin, and 50 µg/mL uridine.

### Compound dose-response curves

Cell lines were plated in 96-well plates (Corning 3903) in 100 µL media at the following densities: NCI-237^UTSW^ (2,000 cells/well); TPC-1 (1,000 cells/well) and allowed to adhere overnight. The following day, the media was exchanged and 200 µL fresh media was added to each well. Compounds (MLN4924 (ApexBio B10356), Bortezomib (Selleck S1013), CD437 (Sigma C5865), (R)-GNE-140 (MedChemExpress HY-100742A), and NCGC00420737 (kindly provided by Gordon M. Stott and the NCI Experimental Therapeutics Program) were dispensed in a 3-fold dilution series using a D300e Digital Dispenser (Tecan). Cells were cultured with compound for 72 hours before viability was assessed using an ATP-based luminescent viability assay, CellTiter-Glo (Promega G7570). Plates were cooled to room temperature for 30 minutes, 100 µL growth media was removed, and 50 µL CellTiter-Glo (prepared according to manufacturer’s directions and diluted 1:1 with PBS + 1% Triton-X-100) was added to each well. Plates were incubated for 10 minutes at room temperature, shaking at 120 rpm on an orbital shaker, before data acquisition using a Synergy 2 or Cytation 5 plate reader (BioTek). For all dose-response curves, data were normalized relative to vehicle-only wells and EC_50_ values were determined using Prism (GraphPad), asymmetric (five parameter), least squares curve fitting. Conditions were assayed in duplicate and all experiments were performed at least twice.

### Forward genetic screening in NCI-237^UTSW^

Pools of wild-type and MSH2-null NCI-237^UTSW^ cells were generated by lentiviral transduction of lentiCRISPRv2 constructs with sgRNAs targeting *AAVS1* or *MSH2*, respectively (approximately 50% survival following puromycin selection based on visual estimate). Cells were cultured for approximately 20-25 passages (estimated 30-40 population doublings) before conducting selections. Experiments were performed with NCI-237^UTSW^ cells cultured in DMEM (Sigma D6429) supplemented with 10% FBS, 1% penicillin/streptomycin, 2 mM L-glutamine, and 50 μg/mL uridine.

Seven-day EC_100_ values were determined as follows: cells were plated in duplicate at 5,000 cells/well in 100 μL media in 96-well plates and allowed to adhere overnight; the following day, media was changed and either 200 μL media (DMEM (Sigma D5796)) supplemented with 1 mM sodium pyruvate (for MLN4924 selections) or 0.1 mM sodium pyruvate (for NCGC00420737 selections) was added to respective wells; compound was dispensed in a 1.5-fold dilution series using a D300e Digital Dispenser (Tecan) and cells were cultured for 7 days without a media change; the EC_100_ value was determined by visual inspection of the plate and assessment of viability with CellTiter-Glo as described above.

Selections were performed as follows: 2 x 10^6^ cells per selection dose were plated in 96-well plates (4 plates per selection dose) at approximately 5,200 cells/well in 100 μL media and allowed to adhere overnight. The following day, media was changed and either 200 μL media with 1 mM sodium pyruvate (for MLN4924 selections) or 0.1 mM sodium pyruvate (for NCGC00420737 selections) with the corresponding amount of compound (seven-day EC_100_; EC_100_ x 1.25; or EC_100_ x 1.5) was added to respective wells using a multi-channel pipette. Cells were cultured for 14 days in the presence of compound with a media change (performed as above) after 7 days with compound. After 14 days in the presence of compound, media was changed and cells were cultured in media without compound (sodium pyruvate increased to 1 mM) for 7 days. To identify potential surviving clones, the media was changed and cells were cultured with 0.04 mg/mL resazurin (Acros Organics AC418900050) in phenol red-free growth media (prepared from Sigma D1145) supplemented with 10% FBS, 1% penicillin/streptomycin, 2 mM L-glutamine, 1 mM sodium pyruvate, and 50 μg/mL uridine for 4 hours in a tissue culture incubator (37°C, 5% CO_2_). Fluorescence was measured on a Cytation 5 plate reader (BioTek) with a 540/35 excitation filter and a 590/20 emission filter to identify wells with surviving cells. Plates were subjected to visual inspection to verify clones identified with resazurin and the media was changed to standard growth media (DMEM (Sigma D6429) supplemented with 10% FBS, 1% penicillin/streptomycin, 2 mM L-glutamine, and 50 μg/mL uridine) without resazurin. The median Z-score for viability was determined as: [(0.6745) * ((Individual Well Fluorescence Intensity – Median Fluorescence Intensity of All Wells) / (Median of the Absolute Value of Individual Well Fluorescence Intensity – Median Fluorescence Intensity of All Wells))]. Surviving clones were expanded in standard growth media without compound and subjected to dose-response assays as described above to assess compound sensitivity.

### *LDHA* and *LDHB* sequencing

Cells were plated in 6-well plates in 2 mL media and allowed to adhere overnight. The following day, media was removed, cells were rinsed one time with 3 mL PBS, and 500 µL TRIzol^TM^ (Thermo Fisher 15596026) was added to each well. Plates were incubated at room temperature for approximately 10 minutes before several rounds of pipetting to lyse cells. The resulting lysate was transferred to a 1.5 mL tube and total RNA isolation was performed as described (61). cDNA was prepared using the High-Capacity cDNA Reverse Transcription Kit (Thermo Fisher 4368813) according to the manufacturer’s instructions with 2 µg total RNA as input. *LDHA* and *LDHB* cDNA were PCR amplified and analyzed by Sanger sequencing.

### Lentivirus production

All lentiviruses were produced by co-transfection of HEK 293T/17 cells with plasmid DNA for the respective lentiviral vector and the packaging components psPAX2 and pMD2.G at a 5:3:2 mass ratio (vector:psPAX2:pMD2.G) using TransIT-LT1 (Mirus MIR 2300). Lentiviral supernatants were collected at 48 and 72 hours post-transfection, pooled, and filtered using 0.45 µm PES filters before immediate use or aliquoting and storage at -80°C.

### Lentiviral expression of LDHA, LDHA^I141T^, LDHB, and mClover

Lentiviral vectors for expression of LDHA or LDHB were generated by PCR amplification of the respective cDNAs from plasmids obtained from the Eugene McDermott Center for Human Growth and Development Sequencing Core Human ORF Collection and cloning into pLVX-IRES-Puro (Takara 632183) through Gibson assembly (NEB E2621). A vector for expression of LDHA^I141T^ was generated through a similar method using overlap extension PCR to introduce the LDHA^I141T^ mutation. NCI-237^UTSW^ and TPC-1 cells were transduced by plating 500,000 cells/well in 6-well plates and adding 100, 300, or 1,000 μL of lentiviral supernatant. Approximately 48 hours after exposure to lentivirus, cells were plated directly into media containing 2 μg/mL puromycin and cultured in puromycin-containing media for 7 days. Cell survival was at least 90% for all cell lines based on visual estimate.

### CRISPR-Cas9 knock-in of LDHA^I141T^

An sgRNA and ssODN pair for HDR-mediated introduction of the LDHA^I141T^ coding sequence at the endogenous *LDHA* locus was identified using the Alt-R HDR Design Tool from IDT. The sgRNA was cloned into PX330 and both a wild-type and LDHA^I141T^ ssODN repair template were ordered as an Ultramer^TM^ from IDT. Approximately 3 x 10^6^ NCI-237 cells cultured in DMEM were plated and allowed to adhere overnight. The following day, cells were transfected with 12 μg PX330-sgLDHA-I141 and 1.2 μg pmaxGFP -/+ 20 μL of 10 μM ssODN. Transfections were performed with Lipofectamine^TM^ 3000 (Thermo Fisher L30000) in Opti-MEM^TM^ (Life Technologies 319-85-062) at a ratio of 1 μg DNA : 100μL Opti-MEM + 1 μL Lipofectamine 3000 + 2 μL P3000 reagent. Approximately 72 hours after transfection, cells were collected, replated to 10 cm dishes at 5 x 10^5^ cells/dish, and allowed to adhere overnight. The following day, the media was changed and cells were cultured with 10 mL media containing 149, 222, 333, or 500 nM NCGC00420737. Media was changed and fresh compound added at 3 days and 7 days after the start of treatment. After 14 days of treatment, media was changed and cells received fresh media with no compound for 7 days before staining with crystal violet (0.5% crystal violet in 20% methanol/water) or picking individual clones for expansion.

### ATP measurements

NCI-237^UTSW^ wild-type and LDHA^I141T^ clones were plated in 96-well plates (Corning 3903) at 15,000 cells/well in 100 µL media and allowed to adhere overnight. The following day, the media was exchanged and 200 µL fresh media was added to each well. Compounds were dispensed using a D300e Digital Dispenser (Tecan). After 4 hour incubation, plates were cooled to room temperature for 30 minutes, 100 µL growth media was removed, and 50 µL CellTiter-Glo (prepared according to manufacturer’s directions and diluted 1:1 with PBS + 1% Triton-X-100) was added to each well. Plates were incubated for 10min at room temperature, shaking at 120 rpm on an orbital shaker, before data acquisition using a Synergy 2 or Cytation 5 plate reader (BioTek). Data were normalized to vehicle-only wells.

### GC-MS metabolomics

For GC-MS metabolomics during LDH inhibitor treatment, NCI-237^UTSW^ cells were plated in 6-well plates at 300,000 cells/well in 3mL media and allowed to adhere overnight. The following day, glucose-free HPLM was prepared and supplemented with 5 mM [U-^13^C_6_] glucose before sterile filtration. Media was further supplemented with 2% dialyzed FBS, 1% penicillin/streptomycin, and 1X insulin-transferrin-selenium. 1 µM (R)-GNE-140, 40 nM NCGC00420737, or an equivalent volume of DMSO was added to media and mixed thoroughly. Plating media was aspirated, cells were rinsed one time with 2 mL PBS, and 2 mL media containing vehicle or LDH inhibitor was added to respective wells. Cells were incubated at 37°C and 5% CO_2_. Following incubation, cells were removed, media was rapidly aspirated, cells were rinsed one time with 4 mL ice-cold saline (0.9% sodium chloride), and 1 mL dry ice-cold extraction solvent (80:20 methanol (Thermo Fisher A456):water (Thermo Fisher W6500)) was added to cells. Cells were scraped and transferred to 1.5 mL tubes and stored at -80°C overnight. The following day, samples were centrifuged at 18,000x *g* for 10 minutes at 4°C, metabolite extracts were transferred to new tubes, and extracts were dried to completeness using a SpeedVac. Samples were derivatized by incubating with 30 µL 10 mg/mL methoxyamine hydrochloride (Sigma 226904) in anhydrous pyridine (Sigma 270970) for 15 minutes at 70°C followed by incubation with 70 µL TBDMS for 1 hour at 70°C. Derivatized samples were transferred to Agilent GC-MS autosampler vials and data acquisition was performed using an Agilent G2579A MSD coupled to an Agilent 6890 gas chromatogram. Data were analyzed using Agilent MSD Chemstation software and a MatLab script for area-under-the-curve analysis, total ion count (TIC) determination, and natural isotopomer abundance correction.

### Immunoblotting

For cell culture samples, cells were plated and allowed to adhere overnight. After treatment, media was removed, cells were washed 1 time with ice-cold PBS, and cells were lysed using Buffer A (50 mM HEPES pH 7.4, 10 mM KCl, 2 mM MgCl_2_) with 1% SDS and 1:2,000 Benzonase® nuclease (Sigma E1014). Cell debris was removed by centrifugation at 16,000x *g* for 5 minutes at room temperature and protein concentration was measured using A_280_ values determined by Cytation 5 (BioTek). Lysates were normalized in Buffer A with 1% SDS and prepared with 6X Laemmli sample buffer containing 100 mM 2-mercaptoethanol (Sigma M6250). Lysates were separated on 4-12% Bolt^TM^ Bis-Tris gels (Invitrogen) using MES-SDS running buffer (Invitrogen B0002) and transferred to PVDF membranes using standard Towbin transfer buffer. After transfer, membranes were stained with 0.1% Ponceau S and stained membrane pictures were captured with a camera. Membranes were rinsed with TBS-0.1% Tween-20 (TBS-T) and blocked with 5% non-fat dry milk in TBS-T for 1 hour at room temperature. Membranes were rinsed with TBS-T and incubated with primary antibodies overnight at 4°C. The following day, membranes were rinsed with TBS-T and incubated with secondary antibodies for 40 minutes at room temperature. Membranes were rinsed with TBS-T, incubated with Clarity Western ECL Substrate (BioRad 1705061) and exposed to X-ray film. Primary antibodies were diluted in 5% bovine serum albumin/TBS-T and used at the following concentrations: anti-MSH2, 1:2,000 (Cell Signaling 2017); anti-LDHA, 1:5,000 (Santa Cruz sc-137243); anti-LDHB, 1:10,000 (R&D Systems MAB9205); anti-GFP, 1:5,000 (Cell Signaling 2965).

### RNA extraction and quantitative PCR (qPCR)

Cells were plated in 6-well plates in 2 mL media and allowed to adhere overnight. The following day, media was removed, cells were rinsed one time with 3 mL PBS, and 500 µL TRIzol^TM^ (Thermo Fisher 15596026) was added to each well. Plates were incubated at room temperature for approximately 10 minutes before several rounds of pipetting to lyse cells. The resulting lysate was transferred to a 1.5 mL tube and total RNA isolation was performed as described (61). cDNA was prepared using the High-Capacity cDNA Reverse Transcription Kit (Thermo Fisher 4368813) according to the manufacturer’s instructions with 2 µg total RNA as input. qPCR was performed in 384-well plates with the following reaction conditions: 5 µL diluted cDNA (1:200 dilution), 5 µL *Power* SYBR^TM^ Green PCR Master Mix (Thermo Fisher 4368577), and 0.25 µL primer mix (1:1:2, 100 µM For primer:100 µM Rev primer:water). Samples were run on a QuantStudio^TM^ 5 system in 384-well plates (Thermo Fisher 4309849). Fold-change in expression values were calculated relative to wild-type NCI-237^UTSW^ using the ΔΔC_t_ method with *TBP* as an internal control.

### Animal studies and NCGC00420737 animal dosing

NCI-237^UTSW^ cell line xenografts were formed by injecting 2 x 10^6^ cells subcutaneously in the flanks of NOD.Cg-*Prkdc^scid^ Il2rg^tm1Wjl^*/SzJ mice (NSG^TM^; Jackson Laboratories strain code 005557). Once tumors were large enough (approximately 500 – 1,000 mm^3^), animals were sacrificed, tumors were dissected, and tumor fragments (approximately 5 mm x 5 mm x 5 mm) were implanted subcutaneously in 8-week-old male NOD-*Prkdc^em26Cd52^Il2rg^em26Cd22^*/NjuCrl (NCG; strain code 572) mice with jugular vein catheters (surgical code: JUGVEIN) and transcutaneous one-channel vascular access buttons (accessory code: INSTBUTON1CH) that were purchased from Charles River Laboratories. Animals were group housed at UT Southwestern Medical Center. NCGC00420737 dosing was initiated when tumors reached approximately 150-200 mm^3^. NCGC00420737 was prepared as a 10 mg/mL stock solution that was dissolved in 0.1 N NaOH/PBS, pH was adjusted to approximately 7.4 using 1 N HCl, and the solution was filtered through a 0.22 μm PES filter. Vehicle was prepared using the same method. A fresh stock of NCGC00420737 or vehicle was prepared weekly and kept at 4°C, protected from light. NCGC00420737 was administered via jugular catheter using the transcutaneous vascular access button for cycles of 5 days on, 2 days off. Tumor and weight measurements were performed daily. Tumor volume was calculated according to [volume = ((Length)*(Width)^2^)*3.14)/6] with the ‘Length’ dimension based on measurements along the rostro-caudal axis and the ‘Width’ dimension based on measurements along the dorso-ventral axis. Animals were sacrificed by CO_2_ asphyxiation approximately 1-1.5 hours after receiving a final dose of NCGC00420737 or vehicle. Tumors were dissected, weighed, and snap-frozen in liquid nitrogen.

### Study approval

Animal studies were approved by the UT Southwestern Institutional Animal Care and Use Committee.

## Data availability

The data and unique reagents generated in this study are available upon request from the corresponding author.

## AUTHOR CONTRIBUTIONS

A.R.F. and D.G.M. conceived the study, designed and interpreted experiments, and wrote the manuscript. A.R.F. and F.V. performed experiments. D.G.M. supervised the study.

## ACKNOWLEDGMENTS

The authors thank the Children’s Research Institute Metabolomics Facility for GC-MS metabolite profiling, and Alex Sternisha for assistance with GC-MS data analysis. We thank members of the McFadden lab for helpful discussions and review of the manuscript. D.G.M. was supported by the Cancer Prevention and Research Institute of Texas (RR140884), a Disease-Oriented Scholar Award from UT Southwestern Medical Center, and the Damon Runyon Cancer Research Foundation (102–19). A.R.F. was supported by the NIH Pharmacological Sciences Training Grant (T32-GM007062). NCGC00420737 was developed using funding from the National Cancer Institute, National Institutes of Health under contract HHSN261200800001E, and with funding from the Chemical Biology Consortium, National Cancer Institute Experimental Therapeutics (NExT) Program.

## SUPPLEMENTARY FIGURES AND FIGURE LEGENDS

**Figure S1, related to Figure 1.**
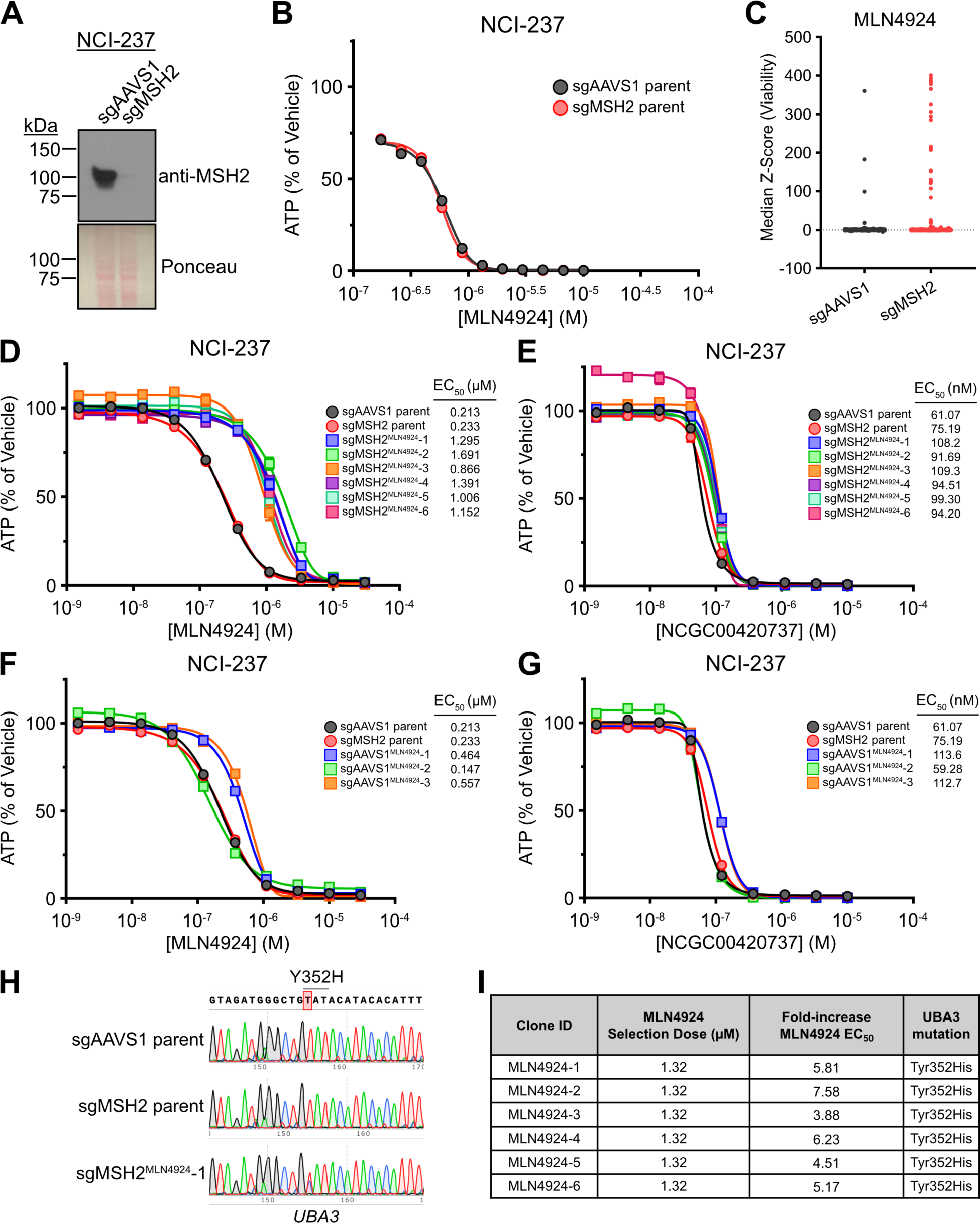
A) Immunoblot for MSH2 in NCI-237^UTSW^. B) Viability assay for NCI-237^UTSW^ cells treated with MLN4924 for 7 days; *n* = 2 replicates. C) Resazurin viability assay for NCI-237^UTSW^ cells treated with 1.32 μM MLN4924 for 14 days; *n* = 384 wells/cell line. Outliers (positive Z-score) indicate viable clones. D) Viability assay for NCI-237^UTSW^ sgMSH2 clones treated with MLN4924 for 3 days; *n* = 2 replicates. E) Viability assay for NCI-237^UTSW^ sgMSH2 clones treated with NCGC00420737 for 3 days; *n* = 2 replicates. F) Viability assay for NCI-237^UTSW^ sgAAVS1 clones treated with MLN4924 for 3 days; *n* = 2 replicates. G) Viability assay for NCI-237^UTSW^ sgAAVS1 clones treated with NCGC00420737 for 3 days; *n* = 2 replicates. H) Sanger sequencing trace for *UBA3* in parental NCI-237^UTSW^ cells and an MLN4924-resistant clone. I) Table of fold-change in EC50 values and *UBA3* mutation status in NCI-237^UTSW^ sgMSH2 MLN4924-resistant clones. Data are plotted as mean ± SEM of the indicated number of replicates. The data displayed in S1D-S1G were generated during the same experiment. The data displayed for sgAAVS1 and sgMSH2 parent in S1D and S1F are the same data. Data displayed for sgAAVS1 and sgMSH2 parent in S1E and S1G are the same data as the data displayed in Fig 1C and 1D.

**Figure S2, related to Figure 1.**
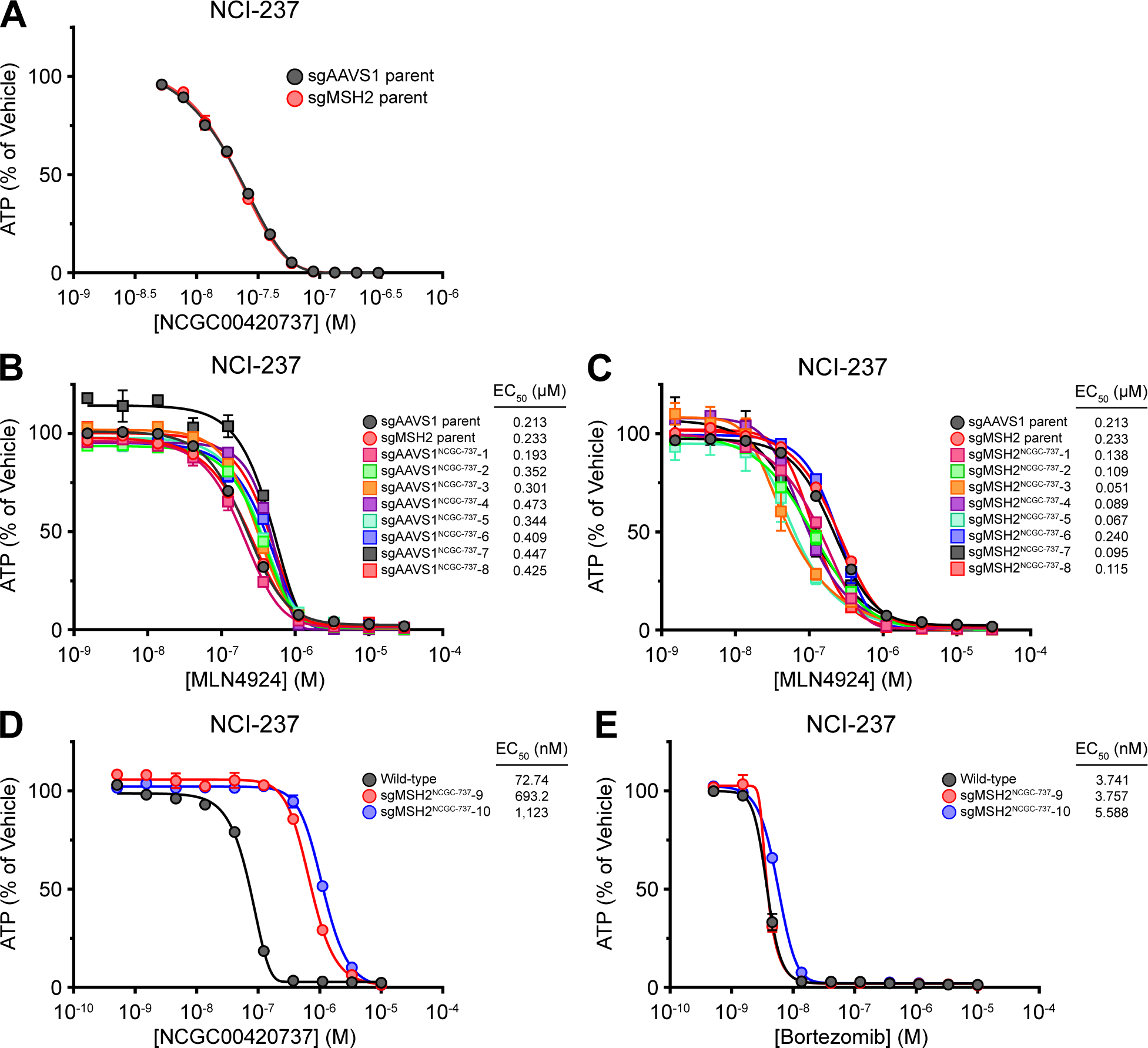
A) Viability assay for NCI-237^UTSW^ cells treated with NCGC00420737 for 7 days; *n* = 2 replicates. B) Viability assay for NCI-237^UTSW^ sgAAVS1 clones treated with MLN4924 for 3 days; *n* = 2 replicates. C) Viability assay for NCI-237^UTSW^ sgMSH2 clones treated with MLN4924 for 3 days; *n* = 2 replicates. D) Viability assay for NCI-237^UTSW^ sgMSH2 clones treated with NCGC00420737 for 3 days; *n* = 2 replicates. E) Viability assay for NCI-237^UTSW^ sgMSH2 clones treated with Bortezomib for 3 days; *n* = 2 replicates. Data are plotted as mean ± SEM of the indicated number of replicates. The data displayed in S2B-S2C were generated during the same experiment as data displayed in Fig 1C-1D and Fig S1D-S1G. The data displayed for sgAAVS1 and sgMSH2 parent in S2B and S2C are the same data, and are the same as the data displayed in Fig S1D and S1F.

**Figure S3, related to Figure 2.**
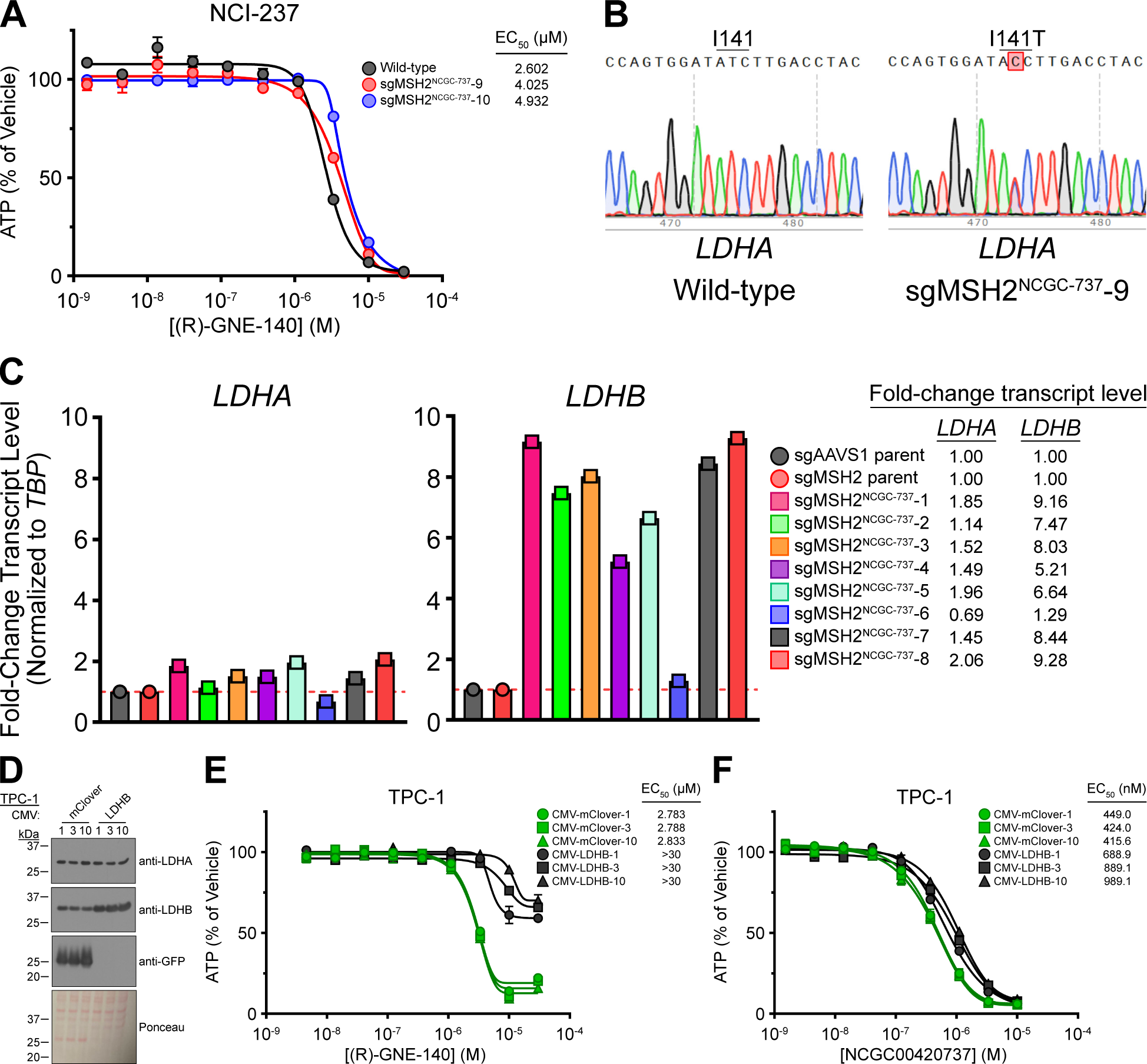
A) Viability assay for NCI-237^UTSW^ sgMSH2 clones treated with (R)-GNE-140 for 3 days; *n* = 2 replicates. B) Sanger sequencing trace of *LDHA* from NCI-237^UTSW^ wild-type and NCGC00420737-resistant (sgMSH2^NCGC-737^-9) cells. C) Quantitative PCR (qPCR) analysis of *LDHA* and *LDHB* expression in indicated NCI-237^UTSW^ cells. D) Immunoblot for LDHA, LDHB, and GFP in TPC-1 cells with lentiviral expression of mClover or LDHB. E) Viability assay for TPC-1 cells with lentiviral expression of indicated constructs treated with (R)-GNE-140 for 3 days; *n* = 2 replicates. F) Viability assay for TPC-1 cells with lentiviral expression of indicated constructs treated with NCGC00420737 for 3 days; *n* = 2 replicates. Data are plotted as mean ± SEM of the indicated number of replicates.

**Figure S4, related to Figure 3.**
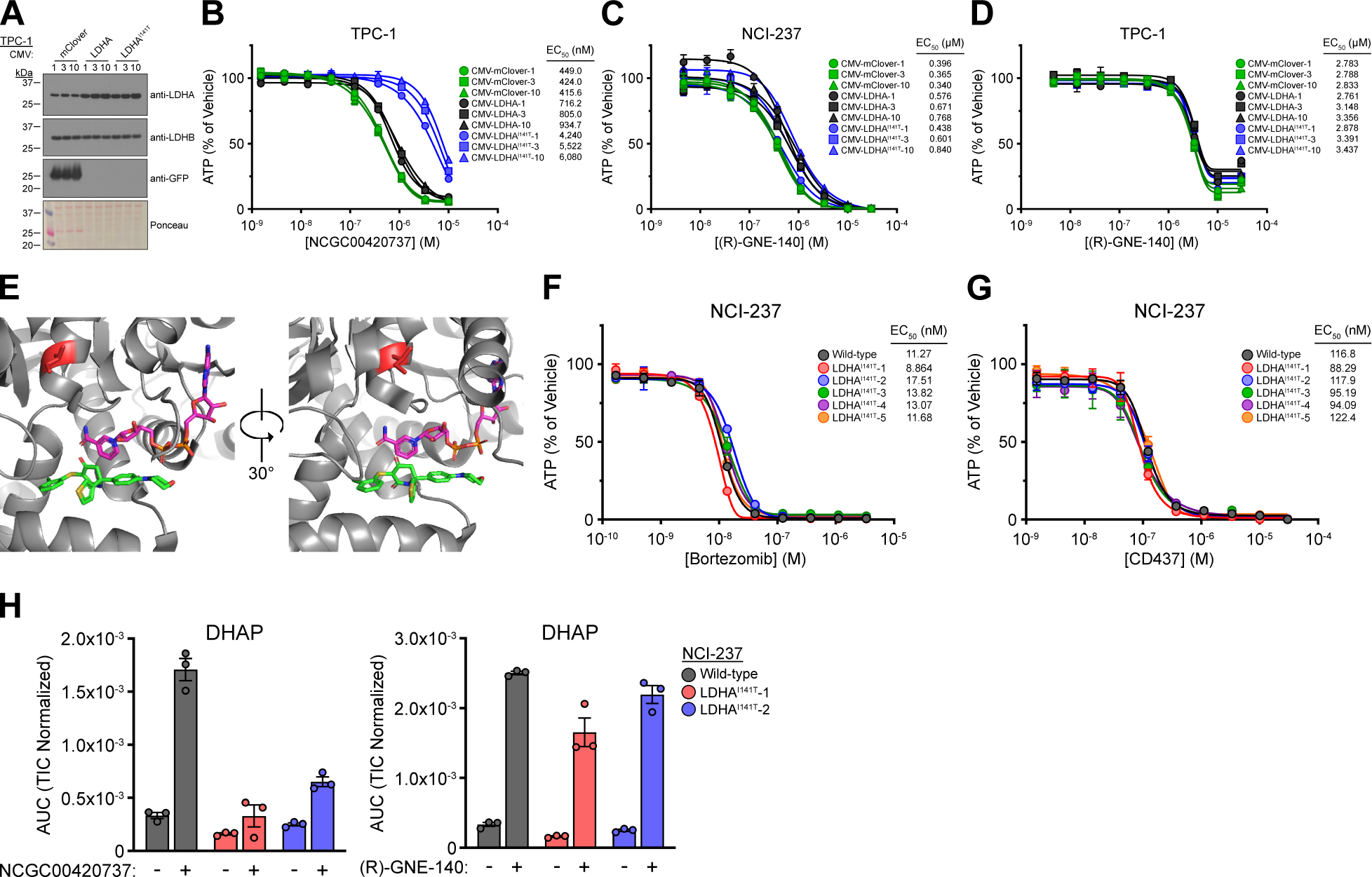
A) Immunoblot for LDHA, LDHB, and GFP in TPC-1 cells with lentiviral expression of mClover, LDHA, or LDHA^I141T^. B) Viability assay for TPC-1 cells with expression of indicated constructs treated with NCGC00420737 for 3 days; *n* = 2 replicates. C) Viability assay for NCI-237^UTSW^ cells with expression of indicated constructs treated with (R)-GNE-140 for 3 days; *n* = 2 replicates. D) Viability assay for TPC-1 cells with expression of indicated constructs treated with (R)-GNE-140 for 3 days; *n* = 2 replicates. E) Co-crystal structure of LDHA and GNE-140. LDHA is colored in gray; Ile141 is colored in red; NAD^+^ is colored in magenta; GNE-140 is colored in green. PDB: 4ZVV. F) Viability assay for indicated NCI-237^UTSW^ cells treated with Bortezomib for 3 days; *n* = 2 replicates. G) Viability assay for indicated NCI-237^UTSW^ cells treated with CD437 for 3 days; *n* = 2 replicates. H) DHAP levels in indicated NCI-237^UTSW^ cells treated with 40 nM NCGC00420737 (left) or 1 µM (R)-GNE-140 (right) for 4 hours; *n* = 3 replicates. Data are plotted as mean ± SEM of indicated number of replicates. The data displayed in S4B-S4D were generated during the same experiment as data displayed in Fig 2E-2F, Fig 3C, and Fig S3E-S3F. The data displayed for CMV-mClover expressing cells in S4B is the same as Fig S3F; data displayed for CMV-mClover expressing cells in S4C is the same as Fig 2E; and the data displayed for CMV-mClover expressing cells in S4D is the same as Fig S3E.

**Figure S5, related to Figure 4.**
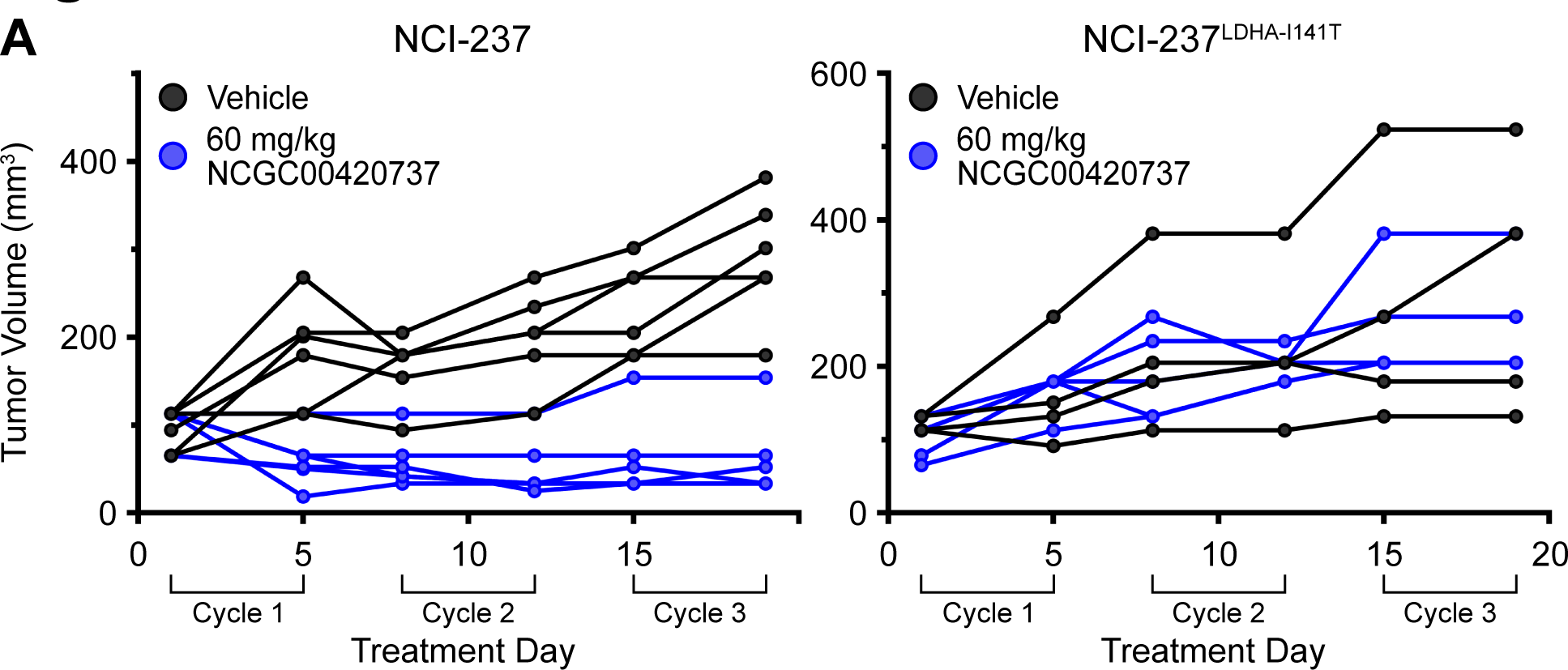
A) Individual tumor volumes from Figure 4.

